# Constitutive and tissue-specific target site preferences of the maize Mutator transposon

**DOI:** 10.64898/2026.05.18.725947

**Authors:** Justin Scherer, Brad Nelms

## Abstract

Transposable elements insert non-randomly into host genomes, generating mutations whose consequences depend on the tissue and developmental stage in which transposition occurs. We analyzed 3.2 million *de novo* Mutator (Mu) insertions across maize leaf, root, pollen, and endosperm to map target site preferences and their tissue specificity at fine-scale resolution. Insertions clustered into 29,408 genomic hotspots near gene 5′ ends, collectively capturing 85% of events while covering just 1.9% of the genome. The vast majority of hotspots were consistently targeted across tissues, indicating that Mu insertion-site preference is largely constitutive rather than tissue-regulated. A minority (∼5%) showed significant tissue-associated variation in Mu activity; this variation was correlated with tissue-specific gene expression, but the relationship was imperfect: some tissue-associated hotspots overlapped broadly expressed genes, while other constitutively active hotspots overlapped genes with tissue-specific expression. These discrepancies indicate that gene expression alone is insufficient to predict Mu insertion-site preference, pointing to additional genomic or chromatin determinants. Together, our results provide a genome-wide, locus-level map of Mu targeting and a quantitative framework for understanding how developmental and genomic context shapes transposon target site preference.

---

Transposable elements (TEs) are mobile DNA sequences that have profoundly shaped eukaryotic genome architecture, with active transposition generating somatic mosaicism (Flasch et al., 2019; Zhang et al., 2020; Movilli et al., 2025; Scherer et al., 2025; Vendrell-Mir et al., 2025), rewiring gene regulation (Bourque et al., 2018; Noshay et al., 2019), and producing heritable variation that contributes to evolution (Stitzer et al., 2021; Liu et al., 2025). Transposon activity is neither uniform across the genome nor across tissues. Germline insertions can be inherited, setting up an evolutionary conflict between transposon proliferation and host fitness (Haig, 2016); this conflict may explain some tissue-specific differences in transposon regulation: the Drosophila P-element transposase is produced selectively in the germline through tissue-specific mRNA splicing (Ghanim et al., 2020), while small RNA pathways restrict transposase activity in both germline and soma (Malone et al., 2009; Slotkin et al., 2009; Matzke & Mosher, 2014). Beyond the germline, many TE families are activated by environmental stress (Pecinka et al., 2010; Dowen et al., 2012; Makarevitch et al., 2015), and TE activity has been implicated in tissue-specific responses to aging (Gorbunova et al., 2021; Siudeja et al., 2021), cancer progression (Rodriguez-Martin et al., 2020), and somaclonal variation in tissue culture (Masuta et al., 2016; Davis et al., 2026).

The consequences of a TE insertion depend sharply on its genomic location. Chromatin features that influence transposon access vary across tissues and developmental stages (Liu et al., 2009; Zhang et al., 2020), and so the same element may generate functionally distinct mutational spectra in different cellular contexts. Consistent with this, stress-activated Copia retrotransposons preferentially insert into loci marked by H2A.Z, a histone variant enriched at environmentally responsive genes (Quadrana et al., 2019; Thieme et al., 2022). Whether such targeting reflects evolved specificity or passive chromatin accessibility remains unclear, connecting to a long-standing debate about the evolutionary significance of non-random mutation rates across the genome (Rosenberg, 2001; Martincorena et al., 2012; Lynch et al., 2016; Xia et al., 2020; Monroe et al., 2022). Either way, understanding the degree to which TE insertion preferences are shaped by tissue and context will be critical to interpreting the impact of TE-induced variation under different conditions and to providing the mutational baseline against which selection signals can be distinguished (Warman et al., 2020).

Capturing TE insertions near the time of transposition, rather than inferring preferences from inherited distributions, allows both somatic and heritable events to be characterized while minimizing the influence of population-level selection. Genome-wide *de novo* TE insertion data have increasingly become available across plant and animal systems (Flasch et al., 2019; Zhang et al., 2020; Movilli et al., 2025; Scherer et al., 2025; Vendrell-Mir et al., 2025; Ambreen et al., 2025), but even large datasets distribute insertions across many millions of genomic base pairs, often with insufficient density to resolve whether specific sites are differentially targeted across tissues or conditions. As a result, prior analyses have largely characterized global genomic trends in aggregate — enrichment near genes, correlation with chromatin marks, chromosomal distribution patterns — without the resolution to identify insertion preferences at individual loci.

In Scherer et al. (2025), we applied MuSeq2 — a sequencing approach with high sensitivity and an estimated false-positive rate of ∼10^-11^ per base pair — to profile *de novo* insertions of the maize Mutator (Mu) transposon across leaf, root, pollen, and endosperm. These four tissues span a broad range of developmental origins, cell division histories, and epigenetic states. With a mean of 73,416 distinct insertion sites per sample and 3.2 million unique sites in total, this dataset provides the combination of density and specificity needed to ask whether individual genomic loci are preferentially targeted in one tissue versus another. Mu is a class II DNA transposon widely used in maize forward genetics (Settles et al., 2007; Marcon et al., 2020) and the founder of the Mutator-like element (MULE) superfamily (Lisch, 2015), with a well-characterized preference for inserting near gene promoters and 5′ UTRs (Zhang et al., 2020). Here, we analyze these data to characterize Mu insertion-site preferences at locus-level resolution across tissues and ask whether genomic and chromatin features explain tissue-associated targeting. Addressing these questions required developing statistical approaches for genome-wide hotspot identification and tissue-association testing that may be applicable to other TE systems.

## RESULTS

### Mu insertion site preferences can be resolved at the locus level in individual tissue samples

We previously obtained MuSeq2 data from 6-17 samples each from leaf, root, pollen, and endosperm (Scherer et al., 2025). In this dataset, individual Mu TE insertions were frequently supported by multiple reads across the TE-genome junction and by reads spanning both the left and right TE borders (**Figure 1**, top), separated by a 9 bp target site duplication characteristic of Mu (Lisch, 2015). There was a mean of 73,416 *de novo* insertion sites per sample (3.2 million insertions in total), providing sufficient density for even small genomic regions to contain multiple independent insertions. **Figure 1** illustrates this in a 500 bp region with relatively high Mu activity (516-fold enrichment over the genomic average): a mean of 9.3 *de novo* insertion sites was detected per Mu-active sample, with no such insertions in Mu-inactive control plants. Although Mu had a statistical tendency to insert at sites with a palindromic CCy<u>CyCnsnGrGr</u>GG motif (target site duplication underlined; **Figure S1**), insertion sites were generally spread throughout the window rather than being dominated by a few locations. For example, across all tissue samples, there were 108 unique insertion locations identified in the 500 bp region shown in **Figure 1** alone.

**Figure 1.**
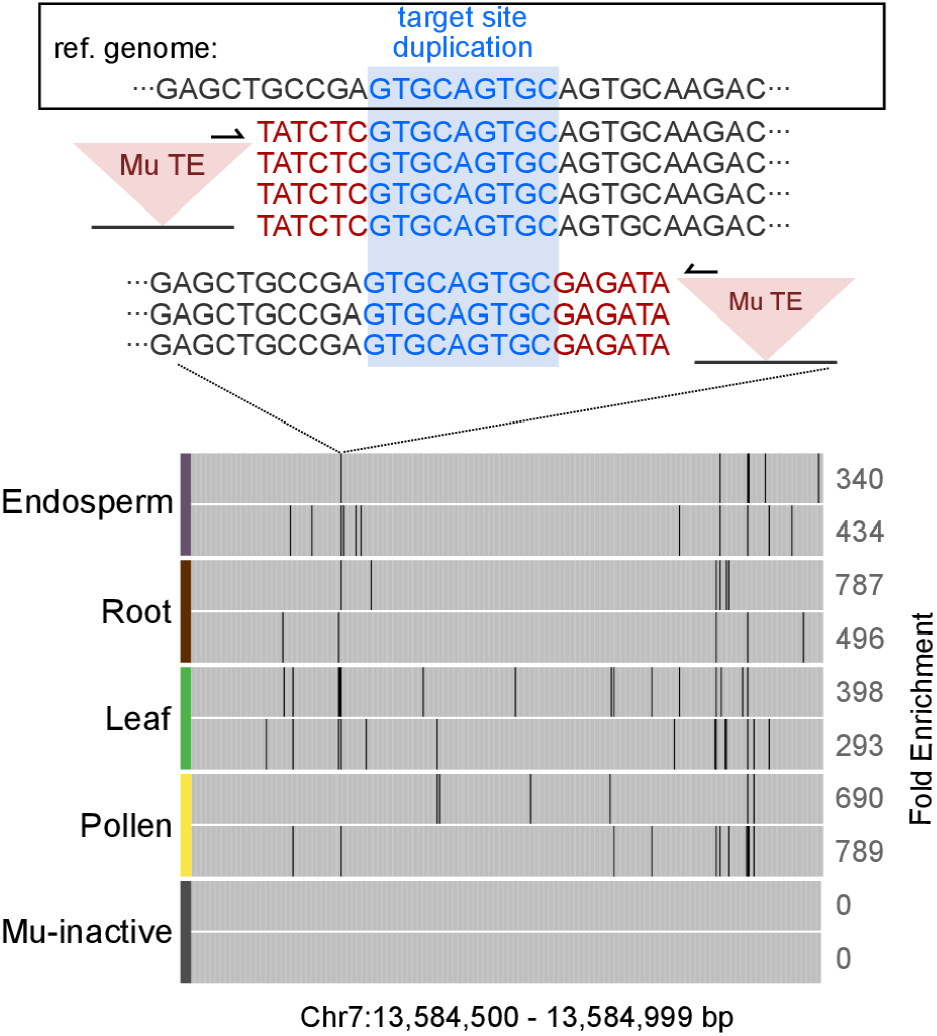
High-confidence detection of *de novo* Mu insertions at individual genomic loci. Top, reads spanning a single Mu insertion site in one endosperm sample; four unique molecules span the right Mu border and three span the left border, with the 9 bp target site duplication indicated by blue shading. Bottom, *de novo* Mu insertion sites (black lines) across a 500 bp genomic region, shown for two samples from each of the four Mu-active tissues and two Mu-inactive control leaf samples. No insertions were detected in Mu-inactive controls. Genome-wide reproducibility across samples is shown in **Figure S2**.

To evaluate the reproducibility of Mu insertion preference across the genome, we divided the genome into 500 bp windows and compared the number of Mu insertions per window between samples. The R^2^ across pairwise sample comparisons was 0.56, significantly greater than zero (*p* < 10^-16^; F-test); mean R^2^ further increased with larger window sizes, indicating that a few hundred bp is the lower bound on spatial resolution. These data therefore provide sufficient resolution to ask whether Mu insertion-site preference varies between tissues at the scale of individual genes.

### *De novo* Mu insertions cluster in hotspots at gene 5’ ends

*De novo* Mu insertions were concentrated in small 0.5-2 kb regions near the 5’ end of genes, consistent with established Mu behavior (Zhang et al., 2020; Lisch, 2015). Mu activity was reproducible between biological replicates, with regions of high enrichment repeatedly observed in different plants (**Figure 2**). Strikingly, the vast majority of these high-activity regions were consistent across all four tissue types – though a subset showed strong tissue-specific differences (**Figure 2**, boxed region) that we characterize in the following section.

**Figure 2.**
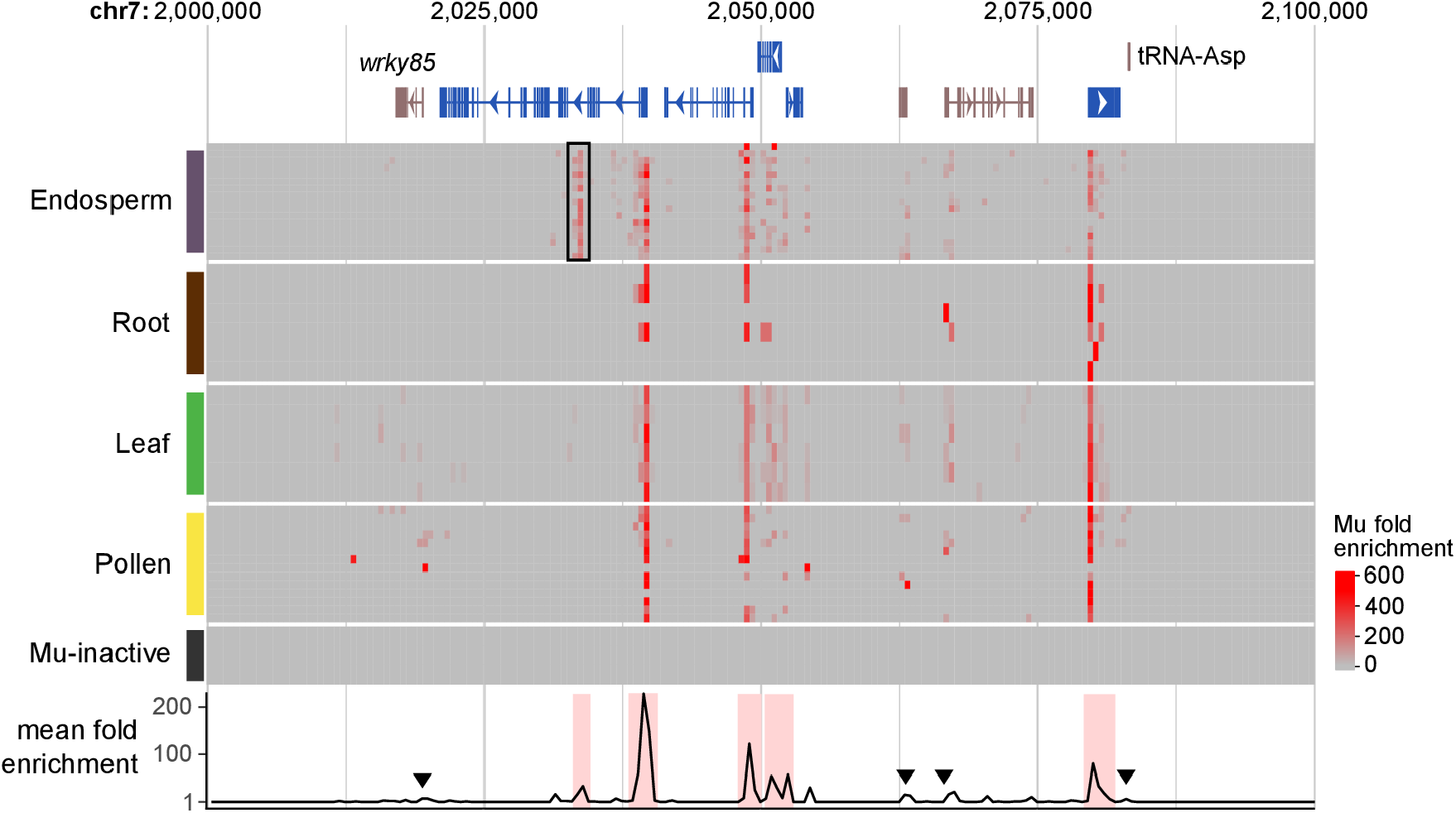
*De novo* Mu insertions are concentrated in hotspots at gene 5′ ends. A 100 kb region of chromosome 7 is shown. Top, gene annotations; blue, genes overlapping a Mu hotspot at their 5′ end; brown, genes without a hotspot. Middle, heatmap of Mu insertion activity across all samples (rows); black box, a hotspot with preferential activity in endosperm samples. Bottom, rolling average Mu fold enrichment across all Mu-active samples in 500 bp windows; red shading, statistically significant Mu hotspots (FDR ≤ 0.01); arrowheads, gene 5′ ends without a called hotspot.

To systematically identify loci with elevated Mu activity (‘hotspots’), we adapted a peak-calling approach analogous to those used for ATAC- and ChIP-seq (Zhang et al., 2008). In total, we found 29,408 Mu hotspots (FDR ≤ 0.01; permutation test), which together covered 1.8% of the mappable genome and captured 85% of *de novo* Mu insertions. Hotspots had a mean size of 1,395 bp (range: 100–9,847 bp) and contained an average of 55 Mu insertions each. This resolution allows us to assess Mu insertion preferences gene by gene — asking not just whether Mu prefers genic regions on average, but which specific genes are targeted and which are not.

Mu hotspots overlapped an annotated gene in 75% of cases, and 57% of maize genes had at least one hotspot. Genes without a hotspot tended to be smaller ([statistics here]) and often showed some elevation of Mu activity above genomic background (**Figure 2**, arrowheads), suggesting the absence of a hotspot sometimes reflects limited statistical power to call weak hotspots rather than a true absence of Mu targeting. Genes without any enrichment in Mu activity tended to be non-Pol II transcripts, such as tRNAs – although some Pol II genes also had consistently low Mu activity (e.g., *Wrky85*; **Figure 2**). These Pol II genes with low Mu insertion rates despite active transcription represent an interesting class, potentially reflecting chromatin or sequence features that restrict transposase access regardless of transcriptional state. There were also hotspots located far from any annotated gene. Many intergenic hotspots overlap regions with detectable RNA-seq signal (Stelpflug et al., 2016), suggesting that some reflect unannotated or misannotated transcription units, while others may represent genuine intergenic Mu activity at loci not captured by gene models.

### A subset of Mu hotspots shows tissue-associated activity

The vast majority of Mu hotspots showed consistent activity across leaf, root, pollen, and endosperm (**Figure 2**), suggesting that the features directing Mu to specific loci are largely constitutive rather than tissue-regulated. To identify the minority of hotspots with genuine tissue-associated activity, we applied an F-test and found that 5.3% had significant variation between tissues (FDR ≤ 0.01). Tissue-associated hotspots had a median 5.9-fold difference in activity between the most and least active tissue, ranging from those with strong tissue-specific activity (**Figure 3B,C**) to those with modest but reproducible variation.

**Figure 3.**
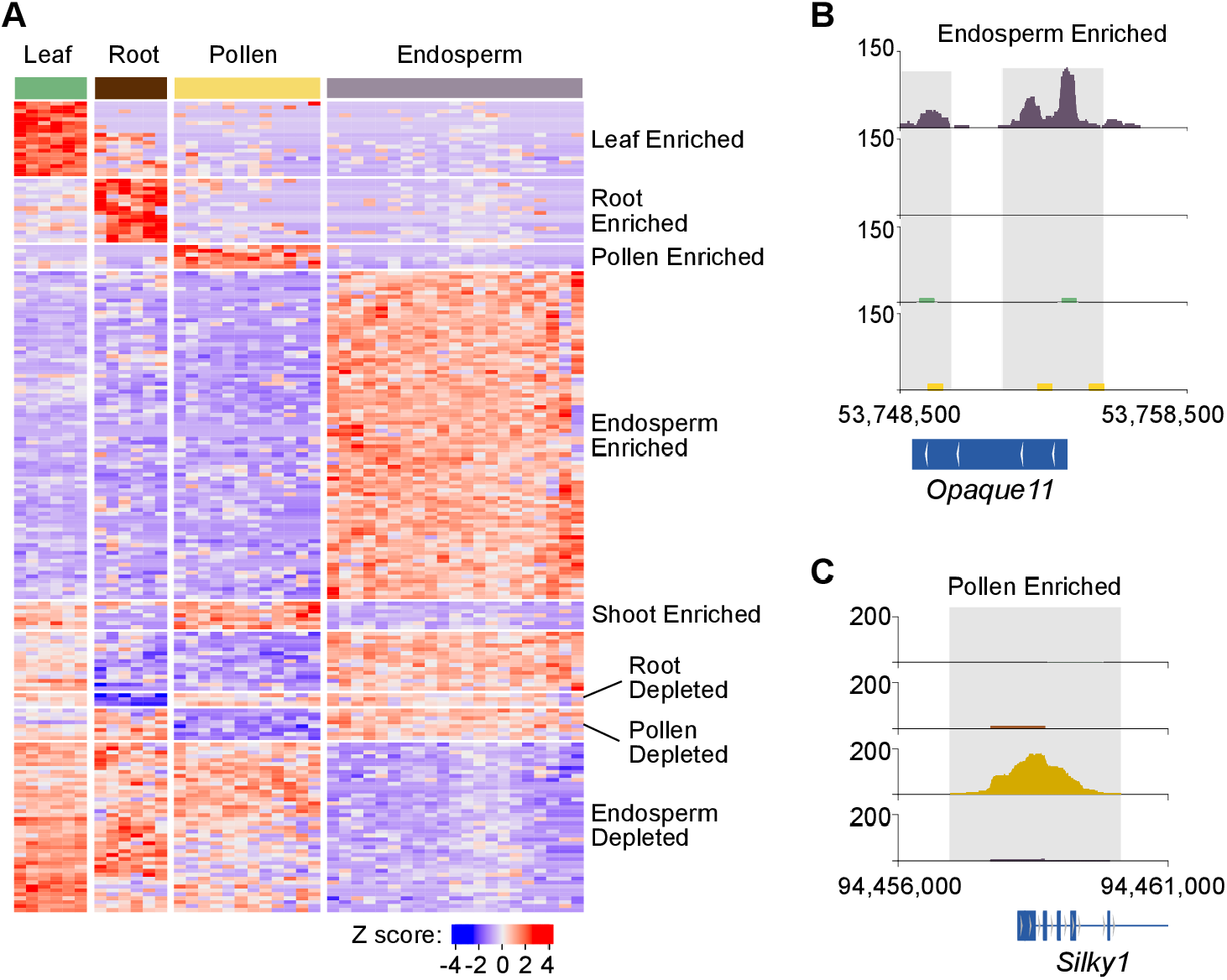
Tissue-associated Mu hotspots. (**A**) Heatmap showing the 200 most significant tissue-associated hotspots (rows) across all samples (columns), with hotspots clustered by pattern of Mu activity across tissues. (**B, C**) Genome browser views of example tissue-associated hotspots; gray shading, hotspot regions. (**B**) Endosperm-enriched hotspot overlapping *Opaque11*. (**C**) Pollen-enriched hotspot overlapping *Silky1*.

We used unsupervised clustering to categorize tissue-associated hotspots based on the pattern of variation in Mu activity. Most tissue-associated hotspots were preferentially active in a single tissue (**Figure 3A**), with the largest cluster (83 hotspots; 41.5%) showing elevated activity in endosperm – a trend that was robust to differences in sample number across tissues. Endosperm has a distinctive chromatin and DNA methylation landscape (Zhang et al., 2013), which may contribute to the prevalence of Mu hotspots with endosperm-specific activity. Clusters with lower activity in one tissue made up a third of tissue-associated hotspots, with endosperm representing the tissue with the most tissue-depleted hotspots.

Two clusters were enriched for Mu in two tissues. One was a shoot-enriched cluster with high activity in both leaf and pollen (**Figure 3A**). This pattern may reflect the shared developmental history of leaf and pollen, both of which are derived from the shoot apical meristem. If Mu insertions at these loci occurred in the meristem or embryo, they would be detected as enriched in both tissues when sampled at maturity. Therefore, the developmental history of a tissue, not just its final state, shapes which loci accumulate insertions.

### Tissue-associated hotspots are partially but imperfectly predicted by tissue-specific expression

Many strong tissue-associated hotspots overlapped genes with matching tissue-specific expression, suggesting that the features directing Mu insertion correlate with those regulating gene expression. For example, a strong endosperm-enriched hotspot overlaps *Opaque11* (**Figure 3B**), an endosperm-expressed bHLH transcription factor whose loss produces small, opaque kernels (Feng et al., 2018). Similarly, a pollen-enriched hotspot overlaps *Silky1* (**Figure 3C**), a class B MADS-box gene expressed in immature tassels and anthers (Stelpflug et al., 2016; Ambrose et al., 2000). Gene expression is known to correlate with Mu activity in aggregate (Zhang et al., 2020), and these examples motivated a more systematic comparison.

We compared Mu fold-enrichment at hotspots that overlapped single genes to tissue-specific expression data (Stelpflug et al., 2016). Mu activity was modestly but significantly correlated with matched tissue expression overall (**Figure 4A**), with endosperm showing the strongest correspondence. Notably, the correlations extended to expression data from developmental stages preceding the sampled tissues: Mu enrichment in leaf correlated with expression in the shoot apical meristem and shoot tip, and Mu enrichment in pollen correlated with expression in the immature tassel and meiotic tassel — consistent with insertion preferences accumulating across a locus’s developmental history, not only in the mature tissue.

**Figure 4.**
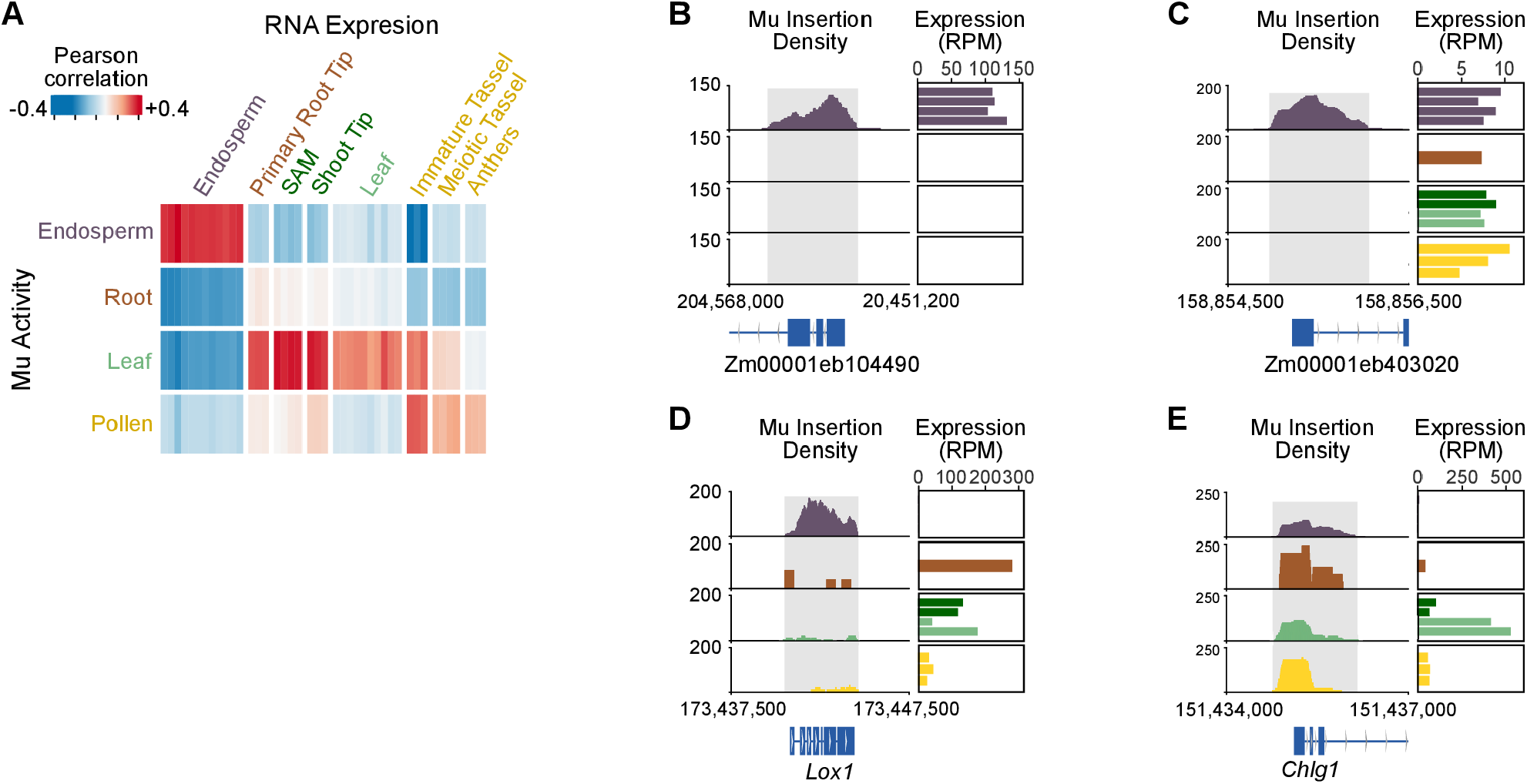
Association between Mu activity and tissue-specific gene expression. (**A**) Heatmap showing Pearson correlation between Mu fold enrichment in each tissue sample (columns) and gene expression in each atlas tissue (rows). Positive correlations indicate that tissues with high Mu activity tend to express the overlapping gene. (**B–E**) Genome browser views of Mu activity (right) and barplots of gene expression across tissues (left) at four loci. (**B**) A concordant example in which endosperm-enriched Mu activity overlaps an endosperm-expressed gene. (**C**) A hotspot with endosperm-enriched Mu activity overlapping a broadly expressed gene. (**D**) A hotspot with endosperm-enriched Mu activity overlapping a gene expressed in all tissues except endosperm. (**E**) A constitutively active hotspot overlapping a leaf-specifically expressed gene (*Chlg1*).

Despite this general trend, tissue-specific expression is neither necessary nor sufficient to predict tissue-associated Mu activity. Overall, from 30% (root) to 80% (pollen) of strong tissue-associated hotspots overlapped genes preferentially expressed in the same tissue or a developmental precursor (e.g., **Figure 4B**). In other cases the correspondence breaks down: for instance, some endosperm-enriched hotspots overlap broadly expressed genes (**Figure 4C**), and others even overlap genes expressed in leaf, root, and pollen but not in endosperm — the tissue where Mu activity is highest (**Figure 4D**). The relationship also fails in the opposite direction: genes with strong tissue-specific expression do not necessarily overlap tissue-associated hotspots. For example, *Chlg1*, a leaf-expressed chlorophyll synthase gene, overlaps a hotspot with uniformly high Mu activity across all four tissues (**Figure 4E**). Together, these results indicate that tissue-specific expression and tissue-associated Mu insertion preference share common determinants at many loci but are clearly dissociable, pointing to additional features beyond transcriptional activity that shape where Mu inserts.

## DISCUSSION

Mu insertion preference is concentrated in hotspots near gene 5′ ends, and the vast majority of these hotspots are active across all four tissues examined. Only ∼5% show significant tissue-associated variation, despite comparing tissues with profoundly different developmental origins, epigenetic landscapes, and gene expression programs. This implies that the primary determinants of Mu insertion preference are constitutive rather than tissue-regulated — a finding that stands in contrast to the expectation that tissue-specific chromatin landscapes would generate tissue-specific insertion spectra.

What contributes to the tissue-specificity of some Mu hotspots? RNA expression provides some predictive power, but is imperfect. Most striking are the ∼5% of endosperm-enriched hotspots that overlie genes expressed in every tissue except endosperm: Mu preferentially inserts at loci in a tissue where the overlapping gene shows no transcriptional activity, which is difficult to reconcile with models in which transcription or associated chromatin state is the proximate driver. Conversely, genes with tissue-specific expression sometimes overlap constitutively targeted hotspots, demonstrating that tissue-specific transcription is not sufficient to generate tissue-specific insertion preference. Together, these cases suggest that Mu target sites are specified by chromatin or DNA features that can meaningfully differ from overall gene expression.

The developmental history of a tissue shapes its Mu insertion profile in ways that transcend its mature state. Pollen-enriched hotspots correlated most strongly with immature tassel rather than mature anther expression, and leaf-enriched hotspots correlated with SAM and shoot tip expression. The shoot-enriched cluster — elevated in both leaf and pollen despite their different mature identities — is most naturally explained by insertions that occurred in shared meristematic or embryonic precursor cells before the two lineages diverged. Because Mu is active throughout development (Scherer et al., 2025), the probability of a given locus accumulating insertions reflects cumulative developmental opportunity — which may help explain why broadly expressed, constitutively accessible genes are more consistently targeted than genes that are only transiently active.

The results here describe insertion preferences for Mu, a DNA transposon with a strong genic bias. Other TE families in maize show different aggregate preferences (Zhang et al., 2020) and may show different degrees of tissue-associated variation — an open question the framework developed here is well-positioned to address as dense *de novo* insertion data become available for other families.

Furthermore, the baseline Mu insertion spectrum may help disentangle insertion preference from selection in existing Mu genetic stocks, where locus-level depletion has been used to infer haploid selection in pollen (Warman et al., 2020) but without knowledge of the underlying insertion preferences. More broadly, the tissue-associated hotspots identified here mark loci where insertion consequences genuinely depend on developmental context, providing a foundation for connecting Mu targeting to its functional and evolutionary consequences in plant genomes.

## Acknowledgments

We thank Bob Schmitz for invaluable discussions. Funding was provided by NIH grant R35GM151237 to B.N.

## Author Contributions

Design and conception, B.N.; Analysis, J.S. and B.N.; Writing, J.S. and B.N.

## Data and materials availability

Upon publication, tissue-associated Mu activity maps will be uploaded to the MaizeGDB genome browser and analysis code will be made available on GitHub. For data use prior to publication, please contact B.N.

## METHODS

### Data acquisition and processing

Mu sequencing data were obtained from GEO accession GSE279993 and remapped to the B73 v5 reference genome (Hufford et al., 2021) to facilitate comparison with B73 gene annotations and publicly available genomic datasets. Genome mapping and Mu insertion site quantification were performed using the same pipeline as described in Scherer et al. (2025), with the exception that reads were mapped to B73 v5 rather than W22. Inherited Mu insertions were identified based on having at least 1,000 molecule counts in both endosperm and a matched sporophytic tissue from the same plant, and were excluded from downstream analyses. A previously published set of 29 historical Mu insertion sites and 19 blacklisted sites (Scherer et al., 2025) were also excluded, resulting in a matrix of *de novo* Mu insertion sites in each tissue sample.

### Hotspot calling

Mu insertion hotspots were identified using a peak-calling approach modeled on the MACS pipeline for ChIP-seq data (Zhang et al., 2008). Insertion site coordinates were first pooled across all samples, retaining each unique genomic position once regardless of how many samples contained it, to create a unified map of insertion sites across the genome.

Hotspot seed regions were identified by counting unique insertion sites in rolling 500 bp windows across the genome. Windows containing fewer than 5 insertions were discarded; under a Poisson null model of uniformly distributed insertions, this threshold corresponds to a one-tailed p-value of approximately 10^-5^, given the observed total number of insertion sites and effective genome size (genome size estimated using deepTools2; Ramírez et al., 2016). Overlapping seed windows were then merged to form candidate hotspots, and the boundaries of each candidate were trimmed to the outermost detected insertion site, so that each hotspot began and ended with an observed insertion.

To assess statistical significance, the −log p-value of each candidate hotspot was calculated under a Poisson distribution. This Poisson p-value serves as a test statistic rather than a nominal p-value, for two reasons. First, candidate hotspots are formed by merging seed regions that already exceeded the Poisson threshold, so their boundaries are defined by regions of elevated insertion density; applying a Poisson test to these merged candidates violates the assumption of a pre-specified window and produces p-values that are anti-conservative. Second, Mu insertion sites are not uniformly distributed across the genome — insertion frequency varies at the chromosomal scale, for example increasing toward chromosome ends — so the uniform Poisson null does not fully capture background variation. Using the Poisson −log p-value as a test statistic, rather than as a nominal p-value, allows us to rank candidates by their degree of local enrichment while correcting for these violations through permutation.

A permutation null distribution was generated by randomly shifting all insertion site coordinates by a distance drawn uniformly between 0 and ±500,000 bp. This shift preserves large-scale chromosomal variation in insertion density while disrupting the local clustering of insertions that defines genuine hotspots. After each permutation, the complete hotspot-calling procedure was repeated to obtain a null distribution of test statistics. The false discovery rate (FDR) for each candidate hotspot was then estimated by comparing the observed test statistic distribution to the permuted null, and hotspots with an FDR ≤ 0.01 were retained.

### Identification of tissue-associated hotspots

Insertion sites within each hotspot were counted per sample and normalized to insertions per million total insertion sites, then converted to fold enrichment relative to the genome-wide expected density (total insertions divided by effective genome size). Fold enrichment values were then log_2_-transformed prior to statistical testing using an empirically determined pseudocount of 10 — log_2_(fold enrichment + 10) — selected to achieve an approximately flat mean-variance relationship across hotspots.

Prior to testing, three classes of hotspots or samples were excluded. Hotspots containing inherited insertions were excluded, as rare Illumina sequencing errors at indel positions can generate artifactual insertion calls adjacent to highly abundant inherited sites. Hotspots with fewer than 50 normalized insertions per million in at least three samples were also excluded, as insufficient Mu activity precludes reliable detection of tissue-associated differences. Finally, two root samples with fewer than 10,000 total insertion sites were excluded, as overall insertion recovery in these samples was too low to reflect biological Mu activity reliably.

To test whether log_2_ fold enrichment varied significantly with tissue type, we applied a permutation F-test to each retained hotspot. Tissue labels were permuted 1,000 times, and the F-statistic from the observed tissue assignments was compared to the permutation null distribution to generate an empirical p-value for each hotspot. False discovery rates were then estimated by comparing the number of hotspots exceeding each p-value threshold in the observed versus permuted data, and hotspots with FDR ≤ 0.01 were designated tissue-associated.

### Genome-browser visualization of Mu insertion-site preference

To visualize Mu insertion-site preference at individual loci, insertion sites from all samples of the same tissue type were pooled and used to compute a rolling average fold enrichment in 500 bp windows. Fold enrichment was calculated as the observed insertion density within each window (insertions per bp) divided by the genome-wide expected density (total insertions divided by effective genome size).

### Comparison to tissue-specific gene expression

RNA-seq data from the Maize Tissue Atlas (Stelpflug et al., 2016) were obtained from the NCBI Sequence Read Archive (accession PRJNA171684) and mapped to the B73 v5 reference genome using HISAT2 with default parameters (Kim et al., 2019). Expression values were normalized to reads per million (RPM); because the comparison of interest is the relative change in expression for a given gene across tissues rather than differences in absolute expression between genes, length normalization does not affect the analysis and RPM was used rather than the more complex RPKM. Expression values were then log_2_ transformed after adding a pseudocount of 10, and Z-transformed across all samples. To ensure a direct one-to-one comparison between hotspot activity and gene expression, only genes overlapping a single tissue-associated hotspot — and hotspots overlapping a single gene — were retained. Pearson correlation was then calculated between the log_2_ Mu fold enrichment and Z-transformed expression values across all pairwise samples.

## Data and code availability

Mu insertion site counts, remapped to the B73 genome, can be downloaded as a processed data file from GEO accession GSE279993. Hotspot locations and fold enrichment across tissues will be made available as supplemental tables and in the MaizeGDB genome browser upon publication. Code for hotspot calling and tissue-association tests will be made available on GitHub upon publication.

## SUPPLEMENTARY FIGURES

**Figure S1.**
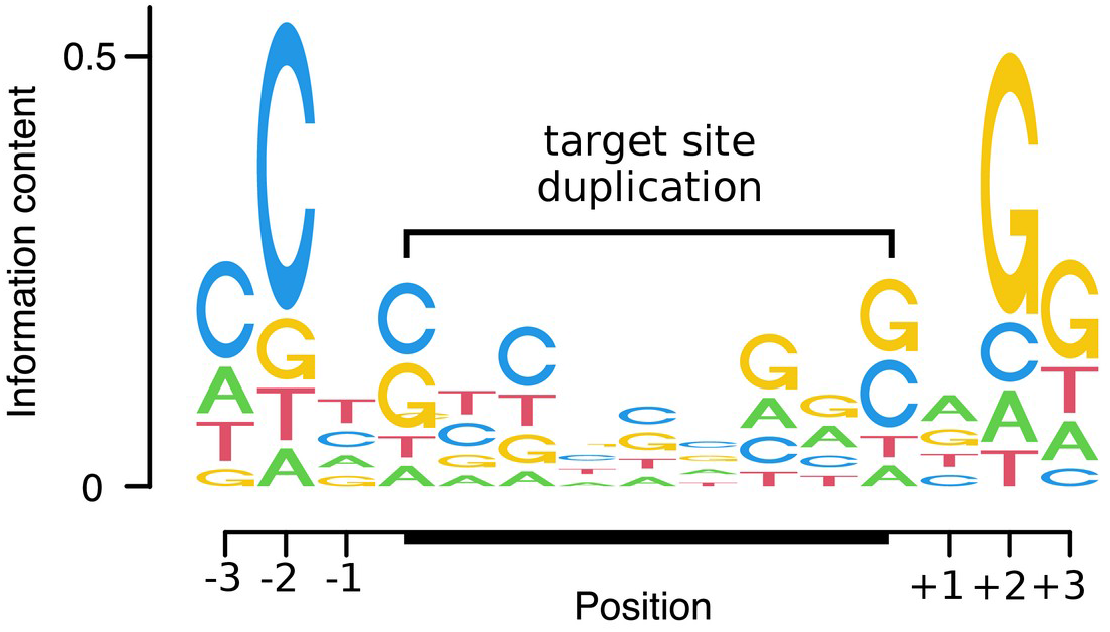
Position weight matrix for genomic sequence at *de novo* Mu insertion sites. For each de novo insertion, the genomic sequence at the precise insertion position is known to single-base resolution from the sequencing reads. The nucleotide frequency at each position surrounding the insertion site was tabulated directly across all de novo insertions and displayed as a sequence logo, with information content (bits) on the y-axis. No motif discovery or peak-calling was required. The central 9 bp correspond to the Mu target site duplication; flanking positions reflect sequence preferences of the transposase beyond the duplication itself.

## REFERENCES

1. Flasch, D. A., Macia, Á., Sánchez, L., Ljungman, M., Heras, S. R., García-Pérez, J. L., Wilson, T. E., & Moran, J. V. (2019). Genome-wide de novo L1 Retrotransposition Connects Endonuclease Activity with Replication. Cell, 177(4), 837-851.e28. 10.1016/j.cell.2019.02.050

2. Zhang, X., Zhao, M., McCarty, D. R., & Lisch, D. (2020). Transposable elements employ distinct integration strategies with respect to transcriptional landscapes in eukaryotic genomes. Nucleic Acids Research, 48(12), 6685–6698. 10.1093/nar/gkaa370

3. Movilli, A., Sushko, S., Rabanal, F. A., & Weigel, D. (2025). Long-read detection of transposable element mobilization in the soma of hypomethylated *Arabidopsis thaliana* individuals. Genome Biology, 26(1). 10.1186/s13059-025-03691-7

4. Scherer, J., Hinczewski, M., & Nelms, B. (2025). Quantitative and sensitive sequencing of somatic mutations induced by a maize transposon. Proceedings of the National Academy of Sciences, 122(32). 10.1073/pnas.2426650122

5. Vendrell-Mir, P., Leduque, B., & Quadrana, L. (2025). Ultra-sensitive detection of transposon insertions across multiple families by transposable element display sequencing. Genome Biology, 26(1). 10.1186/s13059-025-03512-x

6. Bourque, G., Burns, K. H., Gehring, M., Gorbunova, V., Seluanov, A., Hammell, M., Imbeault, M., Izsvák, Z., Levin, H. L., Macfarlan, T. S., Mager, D. L., & Feschotte, C. (2018). Ten things you should know about transposable elements. Genome Biology, 19(1). 10.1186/s13059-018-1577-z

7. Noshay, J. M., Anderson, S. N., Zhou, P., Ji, L., Ricci, W., Lu, Z., Stitzer, M. C., Crisp, P. A., Hirsch, C. N., Zhang, X., Schmitz, R. J., & Springer, N. M. (2019). Monitoring the interplay between transposable element families and DNA methylation in maize. PLOS Genetics, 15(9), e1008291. 10.1371/journal.pgen.1008291

8. Stitzer, M. C., Anderson, S. N., Springer, N. M., & Ross-Ibarra, J. (2021). The genomic ecosystem of transposable elements in maize. PLOS Genetics, 17(10), e1009768. 10.1371/journal.pgen.1009768

9. Liu, B., Munasinghe, M., Fairbanks, R. A., Hirsch, C. N., & Ross-Ibarra, J. (2025). Genomewide selection on transposable elements in maize. bioRxiv. 10.1101/2025.09.16.676665

10. Haig, D. (2016). Transposable elements: Self-seekers of the germline, team-players of the soma. BioEssays, 38(11), 1158–1166. 10.1002/bies.201600125

11. Ghanim, G. E., Rio, D. C., & Teixeira, F. K. (2020). Mechanism and regulation of P element transposition. Open Biology, 10(12). 10.1098/rsob.200244

12. Malone, C. D., Brennecke, J., Dus, M., Stark, A., McCombie, W. R., Sachidanandam, R., & Hannon, G. J. (2009). Specialized piRNA Pathways Act in Germline and Somatic Tissues of the Drosophila Ovary. Cell, 137(3), 522–535. 10.1016/j.cell.2009.03.040

13. Slotkin, R. K., Vaughn, M., Borges, F., Tanurdžić, M., Becker, J. D., Feijó, J. A., & Martienssen, R. A. (2009). Epigenetic Reprogramming and Small RNA Silencing of Transposable Elements in Pollen. Cell, 136(3), 461–472. 10.1016/j.cell.2008.12.038

14. Matzke, M. A., & Mosher, R. A. (2014). RNA-directed DNA methylation: an epigenetic pathway of increasing complexity. Nature Reviews Genetics, 15(6), 394–408. 10.1038/nrg3683

15. Pecinka, A., Dinh, H. Q., Baubec, T., Rosa, M., Lettner, N., & Scheid, O. M. (2010). Epigenetic Regulation of Repetitive Elements Is Attenuated by Prolonged Heat Stress in *Arabidopsis*. The Plant Cell, 22(9), 3118–3129. 10.1105/tpc.110.078493

16. Dowen, R. H., Pelizzola, M., Schmitz, R. J., Lister, R., Dowen, J. M., Nery, J. R., Dixon, J. E., & Ecker, J. R. (2012). Widespread dynamic DNA methylation in response to biotic stress. Proceedings of the National Academy of Sciences, 109(32). 10.1073/pnas.1209329109

17. Makarevitch, I., Waters, A. J., West, P. T., Stitzer, M., Hirsch, C. N., Ross-Ibarra, J., & Springer, N. M. (2015). Transposable Elements Contribute to Activation of Maize Genes in Response to Abiotic Stress. PLoS Genetics, 11(1), e1004915. 10.1371/journal.pgen.1004915

18. Gorbunova, V., Seluanov, A., Mita, P., McKerrow, W., Fenyö, D., Boeke, J. D., Linker, S. B., Gage, F. H., Kreiling, J. A., Petrashen, A. P., Woodham, T. A., Taylor, J. R., Helfand, S. L., & Sedivy, J. M. (2021). The role of retrotransposable elements in ageing and age-associated diseases. Nature, 596(7870), 43–53. 10.1038/s41586-021-03542-y

19. Siudeja, K., van den Beek, M., Riddiford, N., Boumard, B., Wurmser, A., Stefanutti, M., Lameiras, S., & Bardin, A. J. (2021). Unraveling the features of somatic transposition in the Drosophila intestine. The EMBO Journal, 40(9). 10.15252/embj.2020106388

20. Rodriguez-Martin, B., Alvarez, E. G., Baez-Ortega, A., Zamora, J., Supek, F., Demeulemeester, J., Santamarina, M., Ju, Y. S., Temes, J., Garcia-Souto, D., Detering, H., Li, Y., Rodriguez-Castro, J., Dueso-Barroso, A., Bruzos, A. L., Dentro, S. C., Blanco, M. G., Contino, G., Ardeljan, D., … von Mering, C. (2020). Pan-cancer analysis of whole genomes identifies driver rearrangements promoted by LINE-1 retrotransposition. Nature Genetics, 52(3), 306–319. 10.1038/s41588-019-0562-0

21. Masuta, Y., Nozawa, K., Takagi, H., Yaegashi, H., Tanaka, K., Ito, T., Saito, H., Kobayashi, H., Matsunaga, W., Masuda, S., Kato, A., & Ito, H. (2016). Inducible Transposition of a Heat-Activated Retrotransposon in Tissue Culture. Plant and Cell Physiology, pcw202. 10.1093/pcp/pcw202

22. Davis, M. W., Leslie, C. A., Lee, C., Long, E., Meinhold, L., Lorenc, M., Lewis, F., Brown, P. J., & Monroe, G. (2026). Genome degradation in plant tissue culture. Proceedings of the National Academy of Sciences, 123(17). 10.1073/pnas.2530182123

23. Liu, S., Yeh, C.-T., Ji, T., Ying, K., Wu, H., Tang, H. M., Fu, Y., Nettleton, D., & Schnable, P. S. (2009). Mu Transposon Insertion Sites and Meiotic Recombination Events Co-Localize with Epigenetic Marks for Open Chromatin across the Maize Genome. PLoS Genetics, 5(11), e1000733. 10.1371/journal.pgen.1000733

24. Quadrana, L., Etcheverry, M., Gilly, A., Caillieux, E., Madoui, M.-A., Guy, J., Bortolini Silveira, A., Engelen, S., Baillet, V., Wincker, P., Aury, J.-M., & Colot, V. (2019). Transposition favors the generation of large effect mutations that may facilitate rapid adaption. Nature Communications, 10(1). 10.1038/s41467-019-11385-5

25. Thieme, M., Brêchet, A., Bourgeois, Y., Keller, B., Bucher, E., & Roulin, A. C. (2022). Experimentally heat-induced transposition increases drought tolerance in Arabidopsis thaliana. New Phytologist, 236(1), 182–194. 10.1111/nph.18322

26. Rosenberg, S. M. (2001). Evolving responsively: adaptive mutation. Nature Reviews Genetics, 2(7), 504–515. 10.1038/35080556

27. Martincorena, I., Seshasayee, A. S. N., & Luscombe, N. M. (2012). Evidence of non-random mutation rates suggests an evolutionary risk management strategy. Nature, 485(7396), 95–98. 10.1038/nature10995

28. Lynch, M., Ackerman, M. S., Gout, J.-F., Long, H., Sung, W., Thomas, W. K., & Foster, P. L. (2016). Genetic drift, selection and the evolution of the mutation rate. Nature Reviews Genetics, 17(11), 704–714. 10.1038/nrg.2016.104

29. Xia, B., Yan, Y., Baron, M., Wagner, F., Barkley, D., Chiodin, M., Kim, S. Y., Keefe, D. L., Alukal, J. P., Boeke, J. D., & Yanai, I. (2020). Widespread Transcriptional Scanning in the Testis Modulates Gene Evolution Rates. Cell, 180(2), 248-262.e21. 10.1016/j.cell.2019.12.015

30. Monroe, J. G., Srikant, T., Carbonell-Bejerano, P., Becker, C., Lensink, M., Exposito-Alonso, M., Klein, M., Hildebrandt, J., Neumann, M., Kliebenstein, D., Weng, M.-L., Imbert, E., Ågren, J., Rutter, M. T., Fenster, C. B., & Weigel, D. (2022). Mutation bias reflects natural selection in Arabidopsis thaliana. Nature, 602(7895), 101–105. 10.1038/s41586-021-04269-6

31. Warman, C., Panda, K., Vejlupkova, Z., Hokin, S., Unger-Wallace, E., Cole, R. A., Chettoor, A. M., Jiang, D., Vollbrecht, E., Evans, M. M. S., Slotkin, R. K., & Fowler, J. E. (2020). High expression in maize pollen correlates with genetic contributions to pollen fitness as well as with coordinated transcription from neighboring transposable elements. PLOS Genetics, 16(4), e1008462. 10.1371/journal.pgen.1008462

32. Ambreen, H., Leduque, B., Quadrana, L., Slotkin, R. K., Bousios, A., & Nützmann, H.-W. (2025). Somatic mobility of transposons is explosive and shaped by distinct integration biases in Arabidopsis thaliana. bioRxiv. 10.1101/2025.07.14.664700

33. Settles, A. M., Holding, D. R., Tan, B. C., Latshaw, S. P., Liu, J., Suzuki, M., Li, L., O’Brien, B. A., Fajardo, D. S., Wroclawska, E., Tseung, C.-W., Lai, J., Hunter, C. T., III, Avigne, W. T., Baier, J., Messing, J., Hannah, L. C., Koch, K. E., Becraft, P. W., … McCarty, D. R. (2007). Sequence-indexed mutations in maize using the UniformMu transposon-tagging population. BMC Genomics, 8(1). 10.1186/1471-2164-8-116

34. Marcon, C., Altrogge, L., Win, Y. N., Stöcker, T., Gardiner, J. M., Portwood, J. L., II, Opitz, N., Kortz, A., Baldauf, J. A., Hunter, C. T., McCarty, D. R., Koch, K. E., Schoof, H., & Hochholdinger, F. (2020). BonnMu: A Sequence-Indexed Resource of Transposon-Induced Maize Mutations for Functional Genomics Studies. Plant Physiology, 184(2), 620–631. 10.1104/pp.20.00478

35. Lisch, D. (2015). Mutator and MULE Transposons. Microbiology Spectrum, 3(2). 10.1128/microbiolspec.mdna3-0032-2014

36. Zhang, Y., Liu, T., Meyer, C. A., Eeckhoute, J., Johnson, D. S., Bernstein, B. E., Nusbaum, C., Myers, R. M., Brown, M., Li, W., & Liu, X. S. (2008). Model-based Analysis of ChIP-Seq (MACS). Genome Biology, 9(9). 10.1186/gb-2008-9-9-r137

37. Zhang, M., Xie, S., Dong, X., Zhao, X., Zeng, B., Chen, J., Li, H., Yang, W., Zhao, H., Wang, G., Chen, Z., Sun, S., Hauck, A., Jin, W., & Lai, J. (2013). Genome-wide high resolution parental-specific DNA and histone methylation maps uncover patterns of imprinting regulation in maize. Genome Research, 24(1), 167–176. 10.1101/gr.155879.113

38. Stelpflug, S. C., Sekhon, R. S., Vaillancourt, B., Hirsch, C. N., Buell, C. R., de Leon, N., & Kaeppler, S. M. (2016). An Expanded Maize Gene Expression Atlas based on RNA Sequencing and its Use to Explore Root Development. The Plant Genome, 9(1). 10.3835/plantgenome2015.04.0025

39. Feng, F., Qi, W., Lv, Y., Yan, S., Xu, L., Yang, W., Yuan, Y., Chen, Y., Zhao, H., & Song, R. (2018). OPAQUE11 Is a Central Hub of the Regulatory Network for Maize Endosperm Development and Nutrient Metabolism. The Plant Cell, 30(2), 375–396. 10.1105/tpc.17.00616

40. Ambrose, B. A., Lerner, D. R., Ciceri, P., Padilla, C. M., Yanofsky, M. F., & Schmidt, R. J. (2000). Molecular and Genetic Analyses of the Silky1 Gene Reveal Conservation in Floral Organ Specification between Eudicots and Monocots. Molecular Cell, 5(3), 569–579. 10.1016/s1097-2765(00)80450-5

41. Hufford, M. B., Seetharam, A. S., Woodhouse, M. R., Chougule, K. M., Ou, S., Liu, J., Ricci, W. A., Guo, T., Olson, A., Qiu, Y., Della Coletta, R., Tittes, S., Hudson, A. I., Marand, A. P., Wei, S., Lu, Z., Wang, B., Tello-Ruiz, M. K., Piri, R. D., … Dawe, R. K. (2021). De novo assembly, annotation, and comparative analysis of 26 diverse maize genomes. Science, 373(6555), 655–662. 10.1126/science.abg5289

42. Kim, D., Paggi, J. M., Park, C., Bennett, C., & Salzberg, S. L. (2019). Graph-based genome alignment and genotyping with HISAT2 and HISAT-genotype. Nature Biotechnology, 37(8), 907–915. 10.1038/s41587-019-0201-4

43. Ramírez, F., Ryan, D. P., Grüning, B., Bhardwaj, V., Kilpert, F., Richter, A. S., Heyne, S., Dündar, F., & Manke, T. (2016). deepTools2: a next generation web server for deep-sequencing data analysis. Nucleic Acids Research, 44(W1), W160–W165. 10.1093/nar/gkw257

